# A distributed integral control mechanism for the regulation of cholesterol concentration in the human retina

**DOI:** 10.1101/2024.03.14.584346

**Authors:** Ronél Scheepers, Noa L. Levi, Robyn P. Araujo

## Abstract

Tight homeostatic control of cholesterol concentration within the complex tissue microenvironment of the retina is a hallmark of the healthy eye. By contrast, dysregulation of the biochemical mechanisms governing retinal cholesterol homeostasis is thought to be a major contributor to the aetiology and progression of age-related macular degeneration (AMD) in the ageing human eye. Although the signalling mechanisms that contribute to cholesterol homeostasis at the cellular level have been studied extensively, there is currently no systems-level description of the molecular interactions that could explain cholesterol homeostasis at the level of the human retina. Here were provide a comprehensive overview of all currently-known molecular-level interactions involved in the regulation of cholesterol across all major compartments of the human retina, encompassing the retinal pigment epithelium (RPE), the photoreceptor cell layer, the Müller cell layer, and Bruch’s membrane. We develop a detailed chemical reaction network (CRN) of this complex collection of biochemical interactions, comprising seventy-one (71) molecular species, which we show may be partitioned into ten (10) independent subnetworks. These ten subnetworks work together to confer robust homeostasis on thirteen different forms of cholesterol distributed through distinct cellular compartments of the retina. Remarkably, we provide compelling evidence that *three independent* antithetic integral controllers are responsible for the tight regulation of endoplasmic reticulum (ER) cholesterol in retinal cells, and that several *additional independent* mechanisms transfer this homeostatic property to other forms of cholesterol throughout the human retina. Our novel and exquisitely detailed mathematical description of retinal cholesterol regulation provides a framework for considering potential mechanisms of cholesterol dysregulation in the diseased eye, and for the study of potential therapeutic strategies against these pathologies.

## 1 Introduction

In the healthy human eye, vision is accomplished through phototransduction, where photoreceptors in the neural retina receive and transmit visual stimuli to the brain for subsequent processing (Scheepers et al. 2020). The neural retina is a complex sensory tissue found in the posterior part of the eye and lies above a single layer of post–mitotic cuboid cells called the retinal pigment epithelium (RPE) (Fig. 1). The RPE’s apical membrane interacts closely with photoreceptor outer segments via long microvilli, while its basolateral membrane faces Bruch’s membrane (BM), which separates it from the choriocapillaris endothelium. BM consists of five layers; the basal lamina of the RPE, inner collagenous layer (ICL), elastic layer (EL), outer collagenous layer (OCL) and the basal lamina of the choriocapillary endothelium (BL) (Fig 2). Tight junctions between the RPE cells play a crucial role in establishing its apical–to–basolateral polarity and in forming the outer retinal-blood barrier to prevent diffusion of large molecules from the choroid to the retina (Strauss 2005; El-Darzi et al. 2024). Starting from the RPE’s embryonic origin, coordinated maturation requires the cells to adapt to different functional properties of the retina. For example, RPE cells in the macula (the 6 mm diameter region in the center of the retina responsible for central vision) are smaller compared to peripheral RPE cells, have a higher melanin content and are associated with a higher number of photoreceptor cells per RPE cell than in the periphery (Gao and Hollyfield 1992).

**Figure 1.**
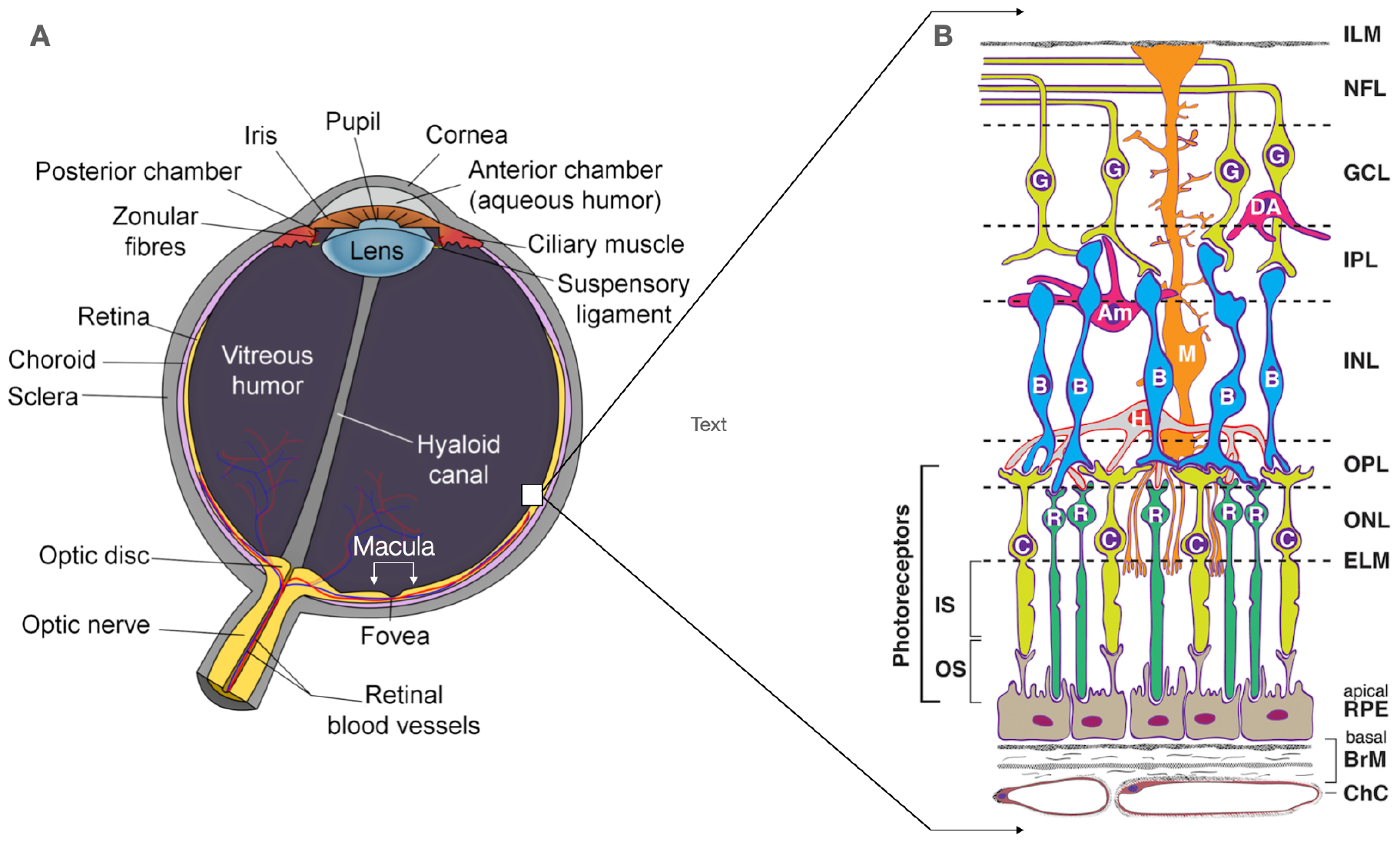
Structure of the retina. **A** Schematic cross–section of the human eye. The neural retina and chororiocappilaris are part of the inner lining, shown in yellow. The macula and fovea (a depression in the macula) are bracketed in white. **B** Chorioretinal cells and layers. Cells: RPE, retinal pigment epithelium; C, cone photoreceptor; R, rod photoreceptor; H, horizontal cell; B, bipolar cell; M, Müller glial cell; Am, amacrine cell; DA, displaced amacrine cell; G, ganglion cell. The area between the foot processes of the Müller cell and the RPE layer forms the subretinal space Ishikawa et al. (2015). Layers: ChC, choriocapillaris; BrM, Bruch’s membrane; ELM, external limiting membrane; ONL, outer nuclear layer; OPL, outer plexiform layer (synapses); INL, inner nuclear layer; IPL, inner plexiform layer; GCL, ganglion cell layer; NFL, nerve fiber layer ; ILM, inner limiting membrane. This image is a derivative of “Schematic diagram of the human eye en.svg” Soerfm (2013). and “Human eye”, taken from (Zheng et al. 2012).

**Figure 2.**
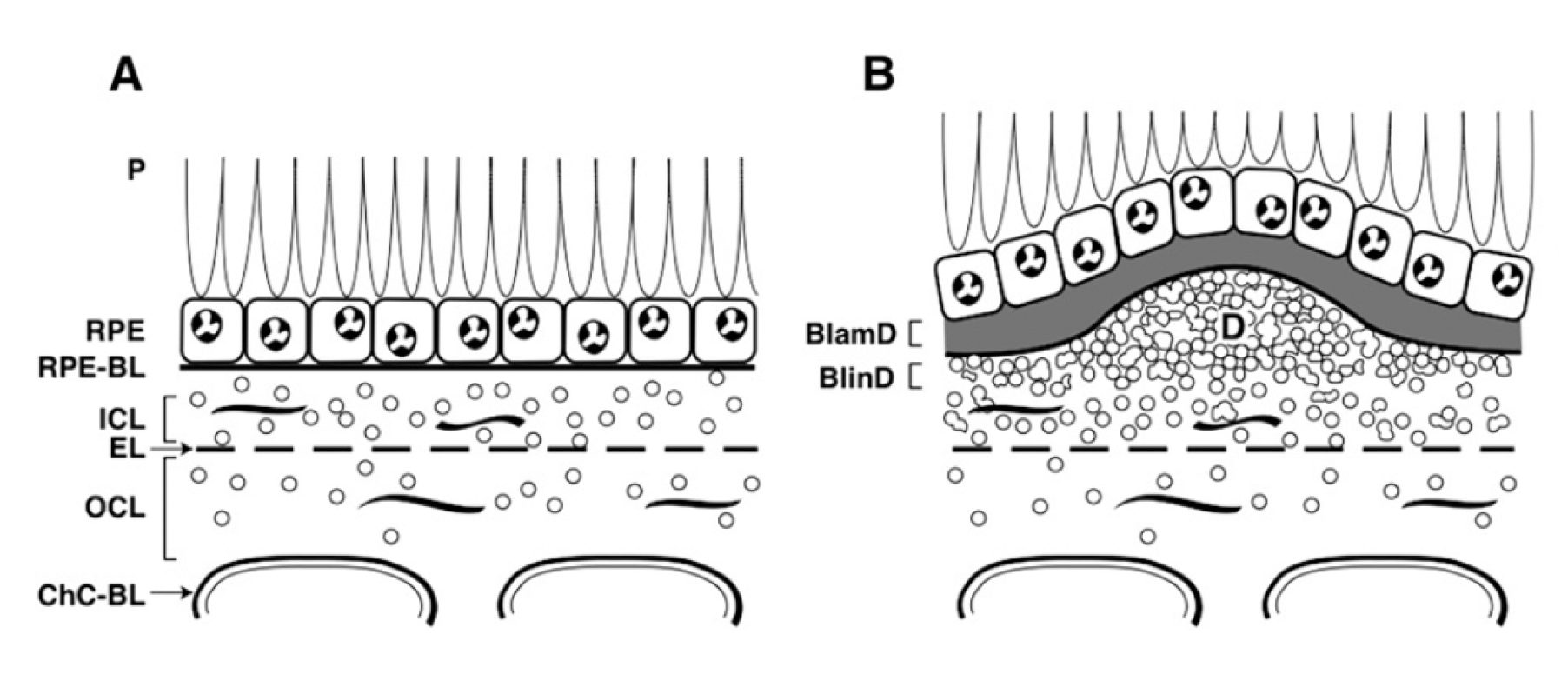
Healthy RPE/BrM versus AMD lesions. Sectional views of the RPE/BrM complex from a healthy eye **A**, compared to an eye with AMD **B**. Small circles represent EC-rich/ApoB lipoproteins, native, and coalesced into drusen (D). **A**: P, photoreceptors; RPE, retinal pigment epithelium; RPE-BL, RPE basal lamina; ICL, inner collagenous layer; EL, elastic layer; OCL, outer collagenous layer; ChC-Bl, basal lamina of choriocapillary endothelium. **B**: BlamD, basal laminar deposit; BlinD, basal linear deposit; D, drusen.(Modified from (Curcio et al. 2010)).

In the diseased state of age–related macular degeneration (AMD), central, acute vision is impaired or lost. Both types of AMD, neovascular AMD (nAMD) and geographic atrophy (GA) share hallmark pathological features – soft drusen and basal linear deposit (BLinD) – constituting an Oil Spill on Bruch’s membrane (BM) (Curcio et al. 2011). Soft drusen and BLinD are two forms of the same extracellular lipid rich material. More specifically, the soft drusen/BLinD deposit forms in the sub retinal pigment epithelium–basal lamina (subRPE–BL) space, an area between the RPE–BL and the inner collagenous layer of BM (See part **B** of Fig 2) (Curcio 2018).

In contrast, pre–BLinD is a layer of three to four rows of densely packed, 60–80nm lipoprotein particles observable in many *healthy* older eyes, in the same space, and first called the Lipid Wall (Ruberti et al. 2003; Curcio et al. 2011; Curcio 2018). Interestingly, it has been shown that these lipoprotein particles first gather in the outer collageonous layer (OCL) of BM in early childhood, and as age increases, these particles gradually fill the subRPE–BL space to form the Lipid Wall (Huang et al. 2007). While various hypotheses exist, it is not clearly understood what age–related changes predispose the progression from constitutively produced individual particles to the pathological state of soft drusen/BLinD development in some individuals.

Significantly, histochemical, biochemical and ultrastructural methods confirmed the presence of unesterified cholesterol and esterified cholesterol in both pre–BLinD and BLinD deposits (Pauleikhoff et al. 1990, 1994; Curcio et al. 2001, 2005). Looking upstream, it has been proposed that these lipoprotein particles are co–translationally assembled in the RPE and deposited into the subRPE–BL space as a way to remove any excess lipids originating from multiple cellular cholesterol metabolism pathways. In particular, excess cholesterol could stem from exogenous uptake of low–density lipoproteins (LDL), high–density lipoproteins (HDL), biosynthesis of cholesterol in the RPE and phagocytosed photoreceptor (PR) outer segment (OS) tips (Curcio et al. 2011; Curcio 2018).

Considering cholesterol’s presence in the pre–BlinD and BLinD deposits in BM and the various intracellular roles cholesterol plays in mammalian retinal cells, a potential link between the homeostatic state of cholesterol in the healthy cell and the consequence of pathology due to disrupted homeostasis is worth investigating. Retinal cholesterol homeostasis entails the interplay between de novo synthesis, uptake, sterol transport, metabolism, and efflux within and between the four main cellular layers of the retina, the choriocapillaris (CC), Bruch’s membrane (BM), the retinal pigment epithelium (RPE) and the light–sensitive cells, also referred to as photoreceptor cells (PR). Defects in these complex processes are associated with several congenital and age-related disorders of the visual system (Pikuleva and Curcio 2014; Curcio 2018).

The fundamental processes in cholesterol metabolism in the vertebrate retina have been studied extensively, particularly under normal physiological conditions. However, how these individual processes are integrated into signalling biochemical networks at intracellular and intercellular levels to achieve cholesterol homeostasis at the tissue level in the retina remains unclear. Only when the many metabolic and signal transduction events involved in cholesterol regulation are fully understood can the potential disruption in signalling pathways, leading to the pathological state, be investigated and elucidated.

In our recent work (Scheepers and Araujo 2023), we have shown that homeostasis is imposed on subcellular cholesterol concentrations in the mammalian cell through a stringent form of regulation known as Robust Perfect Adaptation (RPA) (Araujo and Liotta 2018, 2023c; Khammash 2021). In essence, adaptation signifies a system’s capacity to counteract external disruptions and maintain a crucial internal component, the output, at a constant level. When this output level remains fixed at a specific value, referred to as the ‘setpoint’, independently of the magnitude of disturbances, the system exhibits perfect adaptation. If this stringent adaptation persists despite perturbations to the system’s underlying parameters or structure, it is termed RPA (Gupta and Khammash 2022).

Specifically, we showed that robust homeostasis is achieved through endogenous antithetic integral control, an example of a specific class of molecular control strategies known as *max*RPA (Gupta and Khammash 2022). Having established the foundational molecular mechanisms imposing cholesterol homeostasis in mammalian cells, we seek here to extend our model to investigate the specific molecular mechanisms involved in maintaining cholesterol concentration in the mammalian retina. In doing so, our aim is to identify the detailed molecular mechanisms responsible for conferring RPA on cholesterol concentration in the endoplasmic reticulum (ER) membrane of the retinal pigment epithelium (RPA) cells, the photoreceptors (PR) and the Müller glia (MG) of the retina. Moreover, we seek a mathematical systems-level explanation for: (i) the molecular mechanism by which cells sense a cholesterol concentration in excess of its association capacity with phospholipids in the membrane bilayers, as described in (Scheepers and Araujo 2023), that encompasses the RPE, the PR and the Müller glial cells; and (ii) the molecular mechanisms by which cells maintain a steady state concentration of cholesterol in the pre–BLinD lipoprotein particles.

In Section 2, we comprehensively review the known and theorised molecular interactions and reactions in regulating cholesterol in the four–layer retina. We also include a brief discussion of the critical transcriptional, translational and post–translational signalling events regulating cholesterol identified to exist within and between different subcellular compartments and organelles of the retinal layers. Interested readers are referred to an extensive review of these signalling processes in our recent work here (Scheepers and Araujo 2023). In Section 3, we encode these comprehensive molecular mechanisms into a chemical reaction network (CRN) that induce a set of rate equations under the mass–action assumption that can be analysed computationally to identify RPA–capable species and the molecular mechanisms responsible for RPA in the retina (Section 4).

## 2 The biology of cholesterol in the retina

In the human retina, two types of photoreceptors, rods and cones, function in the phototransduction process, the initial step of vision. There are approximately 120 million rod cells and 6 million cone cells, with the cone cells primarily clustered in the central area known as the macula (Molday and Moritz 2015). To maximise photosensitivity, two opposing processes, disk morphogenesis and shedding, renew the numerous stacked membranous disks that make up these organelles on a daily basis. (Molday and Moritz 2015; Goldberg et al. 2016; Bok 1982).

Recently, independent studies confirmed that disk morphogenesis originate through evaginations of the ciliary plasma membrane (PM) at the proximal end of the photoreceptor cell (Burgoyne et al. 2015; Ding et al. 2015; Volland et al. 2015). In rod photoreceptors, most disks eventually seal off completely to form internalised discrete compartments, while all cone OS disks are continuous with the PM (Goldberg et al. 2016). Photoreceptor cells offset the elongation of their outer segments resulting from continuously adding newly formed disks by actively removing the distal tips of these outer segments. This process, historically referred to as “disk shedding” (Young and Bok 1969), requires engulfment of the OS distal tip by the underlying retinal pigment epithelium cell (RPE) via receptor–mediated phagocytosis. Furthermore, OS shedding is thought to have evolved to remove oxidative lipid species formed in the light–induced environment in the retina. Given that both the photoreceptors and the RPE are post–mitotic, the synchronised renewal of outer segments of photoreceptors is a fundamental retinal mechanism that maintains perpetual retinal homeostasis and function. Moreover, processing phagosomes in tens of thousands of phagocytic events over the lifetime of a human renders the RPE the most active phagocyte in the body (Lieffrig et al. 2023).

Considering that cholesterol is a major lipid and ubiquitous constituent of the PM in mammalian cells, the daily renewal of OS disks requires a prodigious supply of cholesterol to reconstruct both the plasma membranes of the RPE and PR. Specifically, cholesterol forms part of the assembly and function of lipid microdomains in the PM, known as lipid rafts. These dynamic microdomains, rich in unesterified cholesterol and glycosphingolipids, act as a platform for cellular receptors to effectively partition and transport lipids and proteins to the cell membrane through cellular processes such as endocytosis, exocytosis, receptor trafficking and cell signalling (Scheepers and Araujo 2023; Simons and Ikonen 2000; Ouweneel et al. 2020). For these and the many other functions relying on cholesterol (see (Rao and Fliesler 2024)), the total cellular level of cholesterol and its distribution between membranes, and within a given membrane, must be precisely controlled.

Mammalian cells have evolved complex regulatory mechanisms to manage cholesterol levels, ensuring its availability when needed and suppressing cholesterol synthesis and uptake in situations of excess cholesterol (Howe et al. 2016). Cholesterol regulation predominantly occurs in the plasma membrane (PM), endoplasmic reticulum (ER), and mitochondria, where the coordination of *de novo* biosynthesis, uptake, efflux, storage, and chemical modifications supports the maintenance of cellular cholesterol homeostasis (Tabas 2002; Lange and Steck 2016; Luo et al. 2020). It is well–known that the cholesterol regulating machinery resides in the ER, where key proteins respond to abundant cellular cholesterol levels by inhibiting cholesterol–promoting transcription pathways. Specifically, the ER–bound protein SCAP acts as a cholesterol sensor, capturing sterol regulatory element–binding proteins (SREBPs) in the ER when levels are high. Sequestering these transcription factors prevents the activation of genes for cholesterol production and uptake and leads to restored homeostasis. Elevated cholesterol levels furthermore evoke post-translational responses mediated by oxysterols, prompting activation of reverse cholesterol transport pathways and subsequent restoration of lipid equilibrium (Brown et al. 2018; Howe et al. 2016). In our recent work (Scheepers and Araujo 2023), we provide detailed descriptions of these cholesterol regulation pathways.

### 2.1 Cholesterol pathways in the RPE

Considering cholesterol sources in the retina, independent studies by Rao *et al*. (Rao et al. 2018), Biswas *et al*. (Biswas et al. 2017) and Louer *et al*. (Louer et al. 2020) qualitatively demonstrated de novo synthesis of cholesterol in RPE cells. (The interested reader is referred to a summary of key molecules and steps involved in the cholesterol biosynthesis pathway in (Ramachandra Rao and Fliesler 2021)). At the same time, systemic delivery of cholesterol is achieved via the uptake of low–density lipoproteins (LDL) and high–density lipoproteins (HDL) by LDL receptors (LDLR) and scavenger receptors (SR–BI and SR–BII), respectively, which are located in the RPE plasma membrane (Tserentsoodol et al. 2006b; Shen et al. 2018). While the balance of absolute sterol acquisition rates between these two pathways in the RPE is currently unknown, the master regulator of gene expression to realise these pathways is the polytopic ER membrane protein called SCAP (Goldstein et al. 2006). Specifically, Louer *et al*. (Louer et al. 2020) confirmed the expression of the mevalonate pathway gene, HMGCR, in the RPE, where expression of this gene is dependent on the regulated release of the SREBP/SCAP protein complex and its associated transcription factors to the Golgi (Brown et al. 2018). Similarly, the transcription factors required for gene expression of LDLR and SR–BII receptors are regulated through the exact SREBP/SCAP mechanism (Brown et al. 2018; Shen et al. 2018).

Following the daily phagocytosis of the outer segment disk membrane into a nascent phagosome, the phagosome matures by migrating to the basal side of the RPE while undergoing gradual acidification and degradation through phagolysosomal mechanisms to release unesterified cholesterol (UC) (Lakkaraju et al. 2020). Evidence exists that this cholesterol, together with any excess UC in the RPE, could be removed via four efflux pathways: (1) apical efflux via ABSA1 transporters into the subretinal space (Tserentsoodol et al. 2006a; Storti et al. 2017); (2) basal efflux to the circulation (Storti et al. 2017; Ramachandra Rao and Fliesler 2021); (3) localised enzymatic conversion to oxysterols (Mast et al. 2011); and (4) basolateral release as part of a unique APoB-containing lipoprotein into the pre–BlinD layer, from where it enters the circulation via the choriocapillaris, as described in the introduction and shown in Fig 2. (Malek et al. 2003; Curcio et al. 2011).

### 2.2 Cholesterol pathways in the neural retina

A recent review by Rao and Fliesler (Rao and Fliesler 2024) summarises strong evidence from their earlier studies (Fliesler et al. 1993, 1995; Fliesler and Keller 1995) and from other more recent studies (Lin et al. 2015; Clark et al. 2019; Voigt et al. 2020), that the vertebrate retina can synthesise cholesterol *de novo*. Furthermore, based on their review, multiple cell types of the retina most likely have this capacity, including RPE cells, the PR cells and Müller glia.

Significantly, a most recent study involving *in vivo* imaging of the mouse retina to track lipoprotein particles and lipid distribution, provides an explanation of why in situ biosynthesis is the major source of cholesterol for the retina; their data indicates limited trafficking of serum lipoprotein particles (LPP) from the RPE to the neural retina, thus it seems to stimulate *de novo* synthesis of cholesterol in the neural retina (El-Darzi et al. 2024).

Investigating the mechanisms underlying cholesterol biosynthesis and regulation of cholesterol through uptake/efflux pathways in Müller cells, data obtained by (Léger-Charnay et al. 2019) indicates regulation of multiple *de novo* cholesterol synthesis pathway genes, including HMGCR and SREBP2 via 24(S)–hydroxycholesterol, suggesting that Müller cells can efficiently synthesise cholesterol endogenously. Furthermore, their data suggests that Müller cells express the molecular machinery, that is, ABCA1 and ApoE genes, to regulate cholesterol efflux through secretion of ApoE and HDL particles to the subretinal space surrounding neurons.

Given that the interphotoreceptor matrix, a highly organized structure with interconnected domains surrounding cone and rod photoreceptor cells, extend throughout the subretinal space, the cholesterol–carrying HDL particles could be taken up via SR–BI receptors on the PR plasma membrane (Zheng et al. 2012; Ishikawa et al. 2015). Significantly, SR–BI gene expression is regulated by SREBP–1 transcription factors in response to altered intracellular sterol levels (Lopez and McLean 1999). Immunohistochemical analysis of the sterol homeostasis machinery at the protein level reveals that HMGCR, SREBP and SCAP, the proteins of the mevalonate pathway, are expressed in PRs (Zheng et al. 2012). Furthermore, transcript expression profiles of genes encoding the fundamental mevalonate pathway enzymes during retinal development confirms the capacity of PRs to synthesis cholesterol *de novo* (Zheng et al. 2012; Rao and Fliesler 2024)

We summarise all abreviations used in the above descriptions in Table 1 below:

**Table 1.**
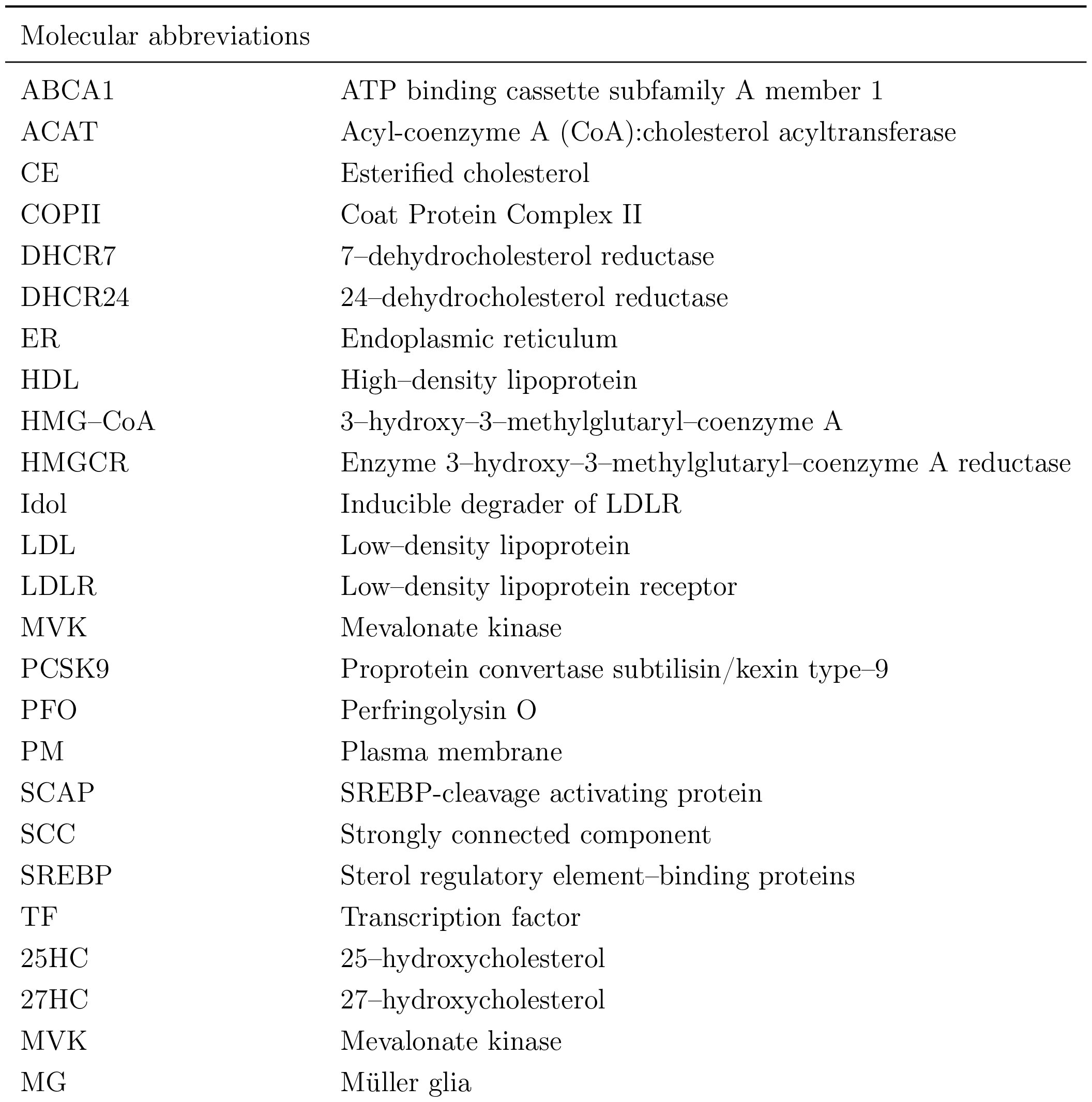

| Molecular abbreviations |  |
| --- | --- |
| ABCA1 | ATP binding cassette subfamily A member 1 |
| ACAT | Acyl-coenzyme A (CoA):cholesterol acyltransferase |
| CE | Esterified cholesterol |
| COPII | Coat Protein Complex II |
| DHCR7 | 7-dehydrocholesterol reductase |
| DHCR24 | 24-dehydrocholesterol reductase |
| ER | Endoplasmic reticulum |
| HDL | High-density lipoprotein |
| HMG-CoA | 3-hydroxy-3-methylglutaryl-coenzyme A |
| HMGCR | Enzyme 3-hydroxy-3-methylglutaryl-coenzyme A reductase |
| Idol | Inducible degrader of LDLR |
| LDL | Low-density lipoprotein |
| LDLR | Low-density lipoprotein receptor |
| MVK | Mevalonate kinase |
| PCSK9 | Proprotein convertase subtilisin/kexin type-9 |
| PFO | Perfringolysin O |
| PM | Plasma membrane |
| SCAP | SREBP-cleavage activating protein |
| SCC | Strongly connected component |
| SREBP | Sterol regulatory element-binding proteins |
| TF | Transcription factor |
| 25HC | 25-hydroxycholesterol |
| 27HC | 27-hydroxycholesterol |
| MVK | Mevalonate kinase |
| MG | Müller glia |

## 3 Methods

### 3.1 CRN representation of the retinal cholesterol regulation network

In this section we represent the biochemical processes discussed in Section 2 as a collection of chemical reactions, thereby generating a Chemical Reaction Network (CRN) for the regulation of cholesterol in the human retina. The forty–five molecular entities involved in this CRN encompass transcription factors, proteins, biochemical intermediates, and macromolecules, as detailed in Table 2. These molecules (‘species’) participate in forty-one chemical reactions, listed in Table 3. Among these species, the extracellular concentrations of cholesterol-carrying lipoproteins (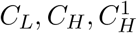 and 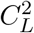) are distinguished as *input molecules* for the system, since these determine the supply of cholesterol to the system (delivered from the bloodstream), and thereby act as the potential disturbances to which the CRN must adapt. In other words, the network must impose RPA on various forms of intracellular cholesterol in response to varying concentrations of these four input molecules.

**Table 2:** List of species and associated symbols in RPE, PR and MG respectively.

| Species | Symbol in RPE | Symbol in PR | Symbol in MG |
| --- | --- | --- | --- |
| SREBP/Scap/Insig complex | $S_{ci}$ | $S_{ci}^1$ | $S_{ci}^2$ |
| Active cholesterol | $C$ | $C_1$ | $C_2$ |
| Cholesterol in ER membrane | $C_e$ | $C_e^1$ | $C_e^2$ |
| Cholesterol in PM | $C_p$ | $C_p^1$ | $C_p^2$ |
| Cholesterol-carrying LDL | $C_L$ | | $C_L^2$ |
| Cholesterol-carrying HDL from circulation | $C_H$ | | |
| Cholesterol-carrying HDL from interphotoreceptor matrix to PR | | $C_H^1$ | |
| LDL receptor (LDLR) | $R$ | | $R_2$ |
| HDL receptor (SR-BI) | $R_b$ | $R_b^1$ | |
| Cholesterol released from receptors | $C_f$ | $C_f^1$ | $C_f^2$ |
| Esterified cholesterol | $E$ | $E_1$ | |
| 3-hydroxy-3-methylglutaryl-coenzyme A (HMG-CoA) | $H$ | $H_1$ | $H_2$ |
| 3-hydroxy-3-methylglutaryl-coenzyme A reductase (HMGCR) | $H_R$ | $H_R^1$ | $H_R^2$ |
| proprotein convertase subtilisin/kexin type-9 (PCSK9) | $P$ | | $P_2$ |
| SREBP transcription factor for LDLR | $S_r$ | | $S_r^2$ |
| SREBP transcription factor for SR-B1 | $S_b$ | $S_b^1$ | |
| SREBP transcription factor for HMGCR | $S_h$ | $S_h^1$ | $S_h^2$ |
| SREBP transcription factor for PCSK9 | $S_p$ | | $S_p^2$ |
| Cholesterol in outer segments of PR | | $C_O$ | |
| Cholesterol in the phagosome | $C_g$ | | |
| ApoB-lipoprotein | $B$ | | |

**Table 3:**
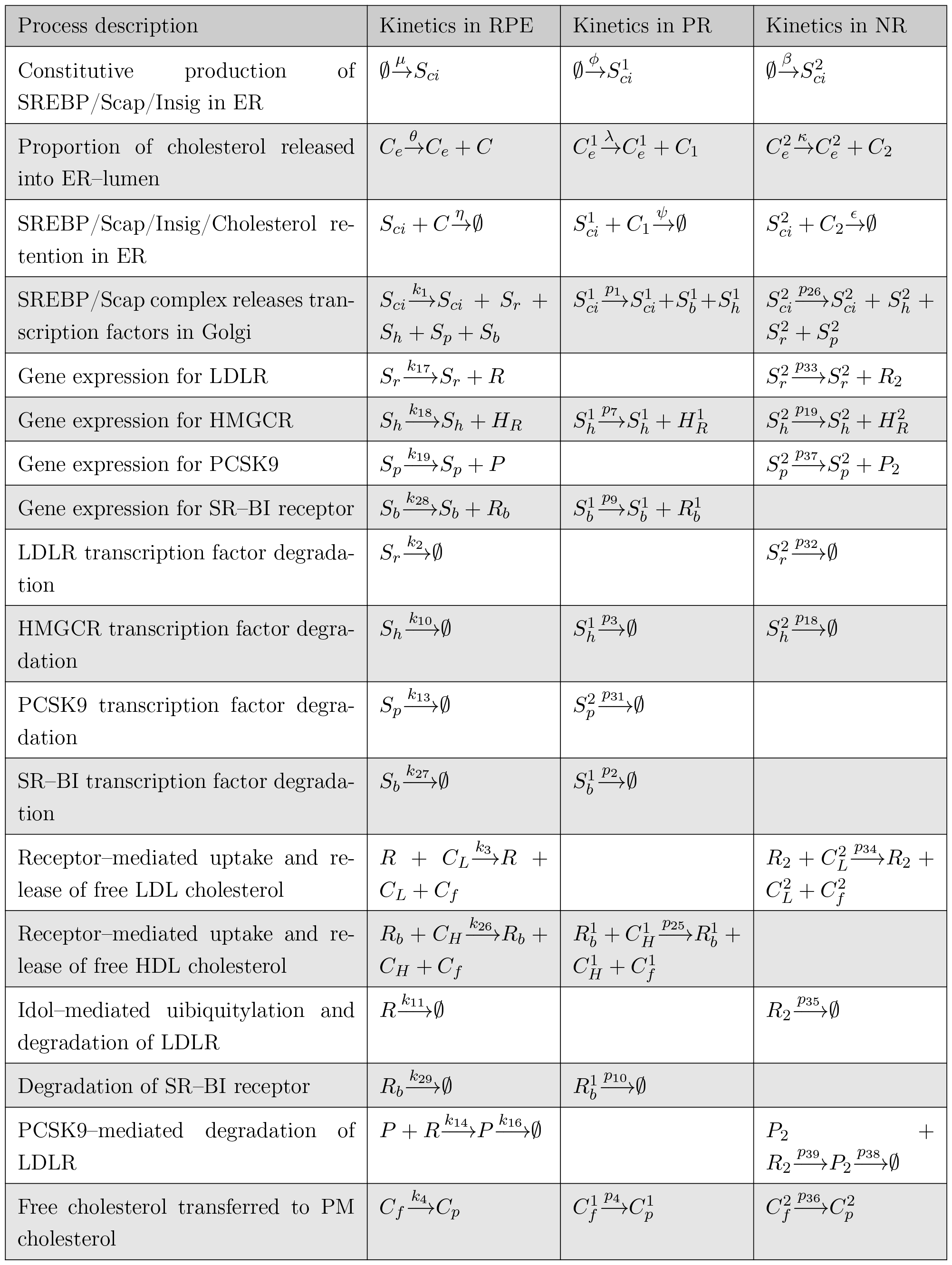
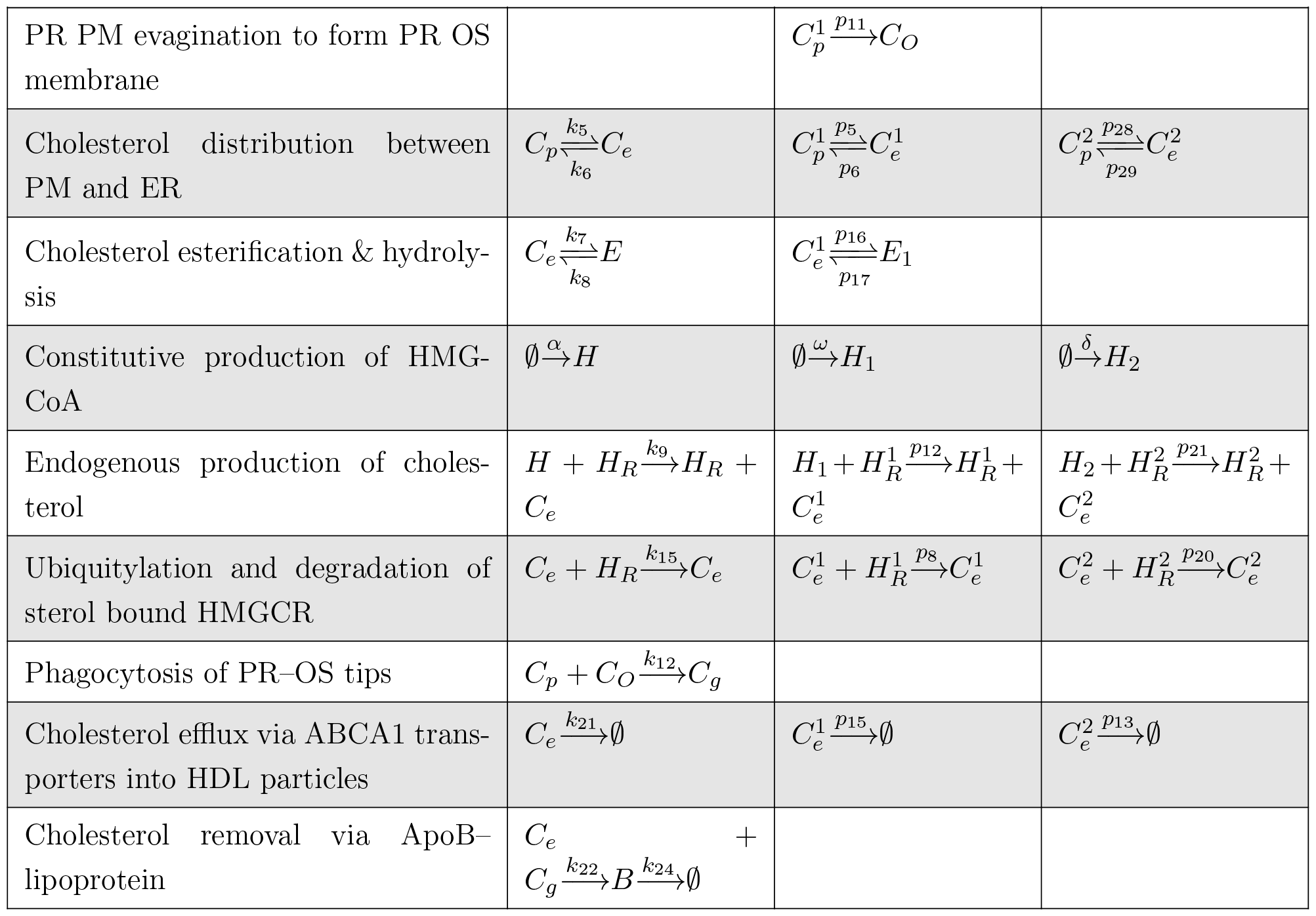
Summary of reactions comprising cellular cholesterol regulation network in the retina.

Using the species nomenclature from Table (2), the biochemical processes captured by the individual reactions of the CRN are as follows, where a superscript or subscript of 1 represents the species in the PR cell, and a superscript or subscript of 2 represents the species in the Müller cell. When referring to the general case, we use the notation *S*^*i*^ or *S*_*i*_:

The constitutive production rate of the SREBP/SCAP/Insig1 complex in the ER is represented by the reaction 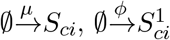 and 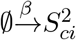 respectively (Brown et al. 2018).

In conditions where cholesterol is depleted, the *S*_*ci*_ complexes are transported to the Golgi for processing. This model does not include the dissociation and degradation of Insig-1. The Golgi’s proteolytic activity facilitates the release of the SREBP transcription factors into the nucleus.

In the RPE, this process is represented by the equation 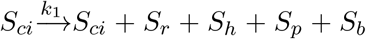, while transcription factor release in the PR is modelled as 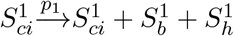 and in the Müller cell 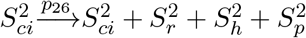 as.

The rate at which the transcription factors *S*_*r*_ and 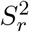 activates transcription of its target genes to produce LDLR protein is captured as 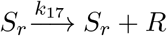 in the RPE and 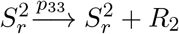 in the Müller cell respectively. Activation of the genes transcribing SR–BI receptor protein production is facilitated by the transcription factors *S*_*b*_ and 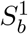, and modelled as 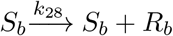 in the RPE and 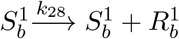 in the PRs respectively.

In a similar manner, *S*_*h*_, 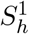 and 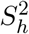 induces the synthesis of HMGCR in the ER of the three retinal layers, represented as 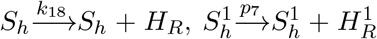 and 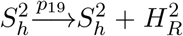 respectively.PCSK9, the enzyme promoting degradation of LDLR, is produced through *S*_*p*_ and 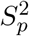 transcription, and represented as 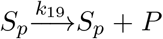 and 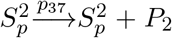 respectively (Xia et al. 2021). The degradation rates of the SREBP transcription factors are denoted as 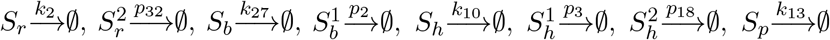 and 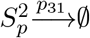 respectively (Sundqvist and Ericsson 2003).

The reactions 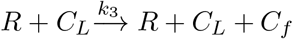 and 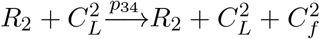 represent the receptor– mediated uptake and subsequent release of unesterified cholesterol through LDLR into endosomes in the RPE and Müller cells respectively. Similarly, the receptor–mediated uptake and release of cholesterol through SR–BI receptors into endosomes in the RPE and PRs are modelled as 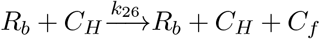 and 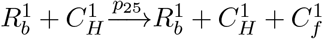 respectively. Here, the conservation of *R*_*i*_ represents the recycling of either LDLR or SR–BI to the surface membrane. For a detailed understanding of this recycling mechanism, readers are directed to the works of (Tindall et al. 2009) and (Pool et al. 2018).

Post-translational regulation of LDLR involves IDOL-mediated degradation, which is succinctly represented by the equations 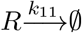 and 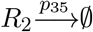, while the binding of PCSK9 to LDLR facilitates the degradation of LDLR as well as its own degradation in the lysosomes. These degradation processes are described by the equations 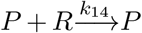 and 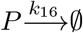 in the RPE and 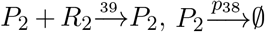 in the Müller cells.

The movement of cholesterol released from the respective receptors 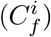 to the plasma membrane (PM) is represented by the rate equations 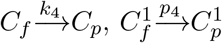 and 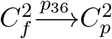. From here, bidirectional fluxes between cholesterol in the PM, 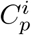, and the ER membrane cholesterol pool, 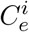, are captured as 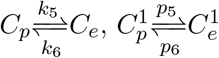 and 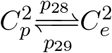 respectively.

A fraction of the cholesterol in the ER is transformed into cholesterol esters at a rate represented by 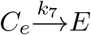 and 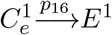, and conversely, a proportion of *E* can be reverted to unesterified ER cholesterol as needed, depicted by the rate 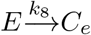 and 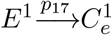 respectively. Given that cholesterol in the neural retina is mostly unesterified under normal conditions (*∼* 87%), modelling the conversion of UC into cholesterol esters in the Müller cells was omitted (Saadane et al. 2016; Bretillon et al. 2008).

Regarding the cholesterol biosynthesis pathways in the RPE, PRs and Müller cells respectively, the constant rate of HMGCoA (*H*_*i*_) production is denoted by the equations 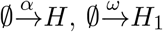 and 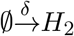.

Following a series of omitted intermediate steps, the end product of the biosynthesis pathway, 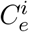, is generated through the activation of HMGCoA by the HMGCR reductase enzyme. This process is represented by the equations 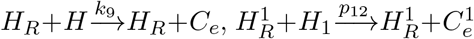 and 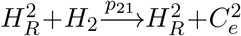 respectively.

The degradation of HMGCR, a key step in cholesterol biosynthesis and regulated by feedback mechanisms, is represented by the equations 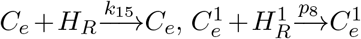 and 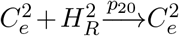.

ER cholesterol molecules, *C*_*i*_, exceeding the phospholipid bilayer’s sequestration capacity – that is, the *active cholesterol molecules* –, are released into the ER lumen of the RPE, PRs and Müller cells, and modelled by 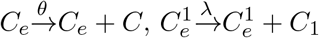 and 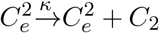 respectively.

Cholesterol molecules in the ER lumen are sensed and captured by the 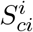 complex, resulting in the formation of a stable 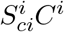 complex within the ER membrane, as represented by 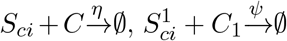, and 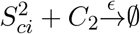. This process prevents the 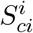 complex from moving to the Golgi and releasing SREBP transcription factors, ultimately decreasing the concentration of cholesterol in the cell.

Excess cholesterol in the various cells are removed via ABCA1 transporters into HDL–like particles to either the choriocapillaris or the subretinal space, and these processes are represented as 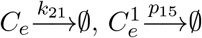 and 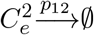 (Curcio et al. 2011).

PR PM evagination to form the PR OS is modelled as 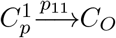. The daily phagocytosis of PR OS membrane tips involves engulfment from the RPE PM, and the reaction equation representing this process is 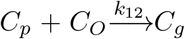, where *C*_*g*_ represents the cholesterol in the phagosome in the RPE. The cholesterol released from the phagosome, together with any excess ER cholesterol are removed to the choriocapillaris basolaterally and this process is modelled as 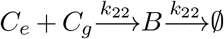.

Figure 3 presents a schematic summary of these interactions, outlining the overall network structure involved in regulating cholesterol in the retina.

**Figure 3.**
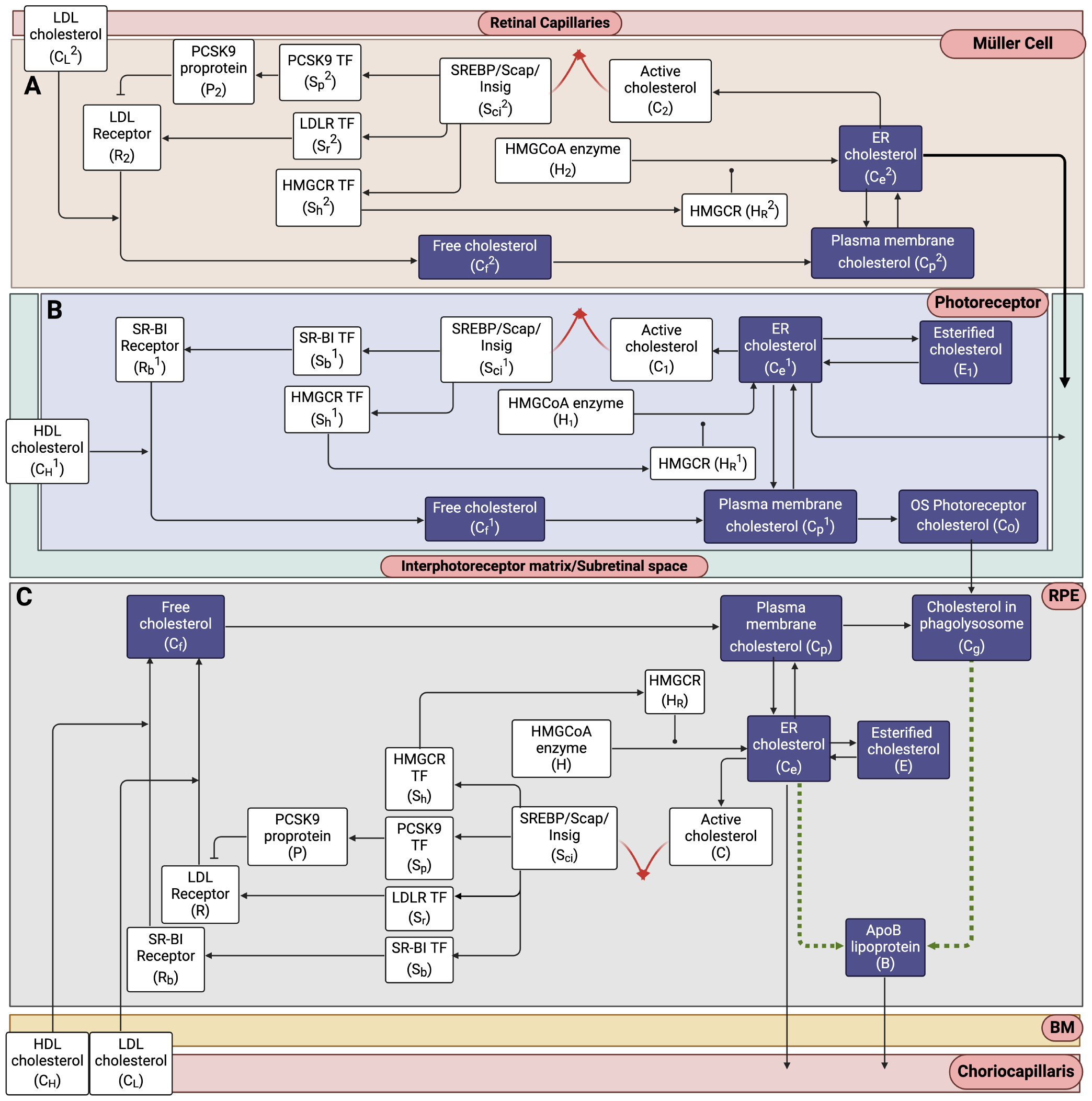
Network schematic for cellular cholesterol regulation in the retina. The presented network diagram captures the interplay between the 41 species of the CRN, unveiling the overarching organizational structure of the cellular cholesterol homeostatic apparatus as a singular Opposer module in **A**, the MG, **B**, the PRs and **C**, the RPE, characterized by its distinctive feedback architecture. Arrows depict flux (activation), blunt ends depict inhibition, and solid circles denote catalysis. Red arrows signify the opposing nodes: irreversible cholesterol sequestration in the 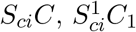 and 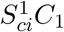 complexes. Dashed dark green arrows represent the proposed assembly of a unique ApoB lipoprotein particle characteristic to the retina. (Created with BioRender.com)

### 3.2 Chemical Reaction Rates

Under the mass-action assumption, the CRN detailed in the preceding section induces the following polynomial dynamical system of forty-one reaction rates, which together capture the dynamical regulation of retinal cholesterol:

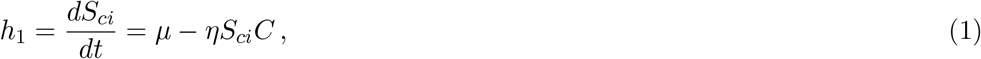

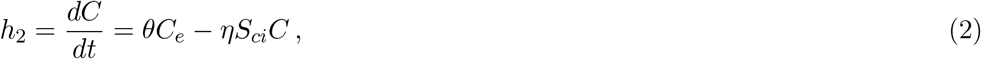

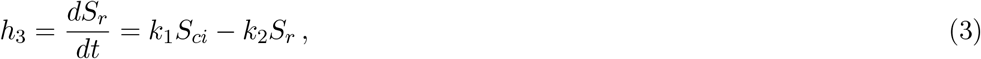

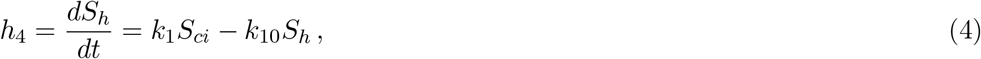

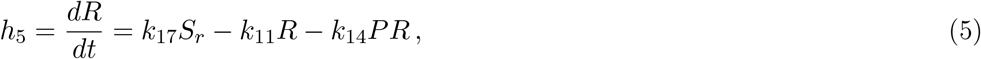

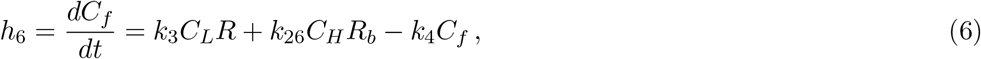

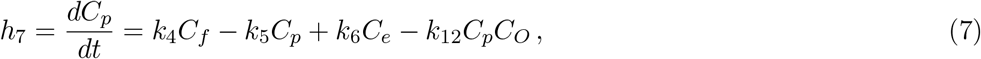

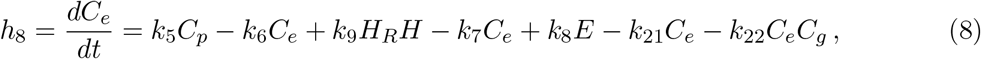

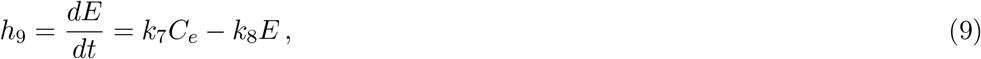

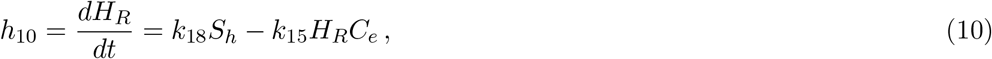

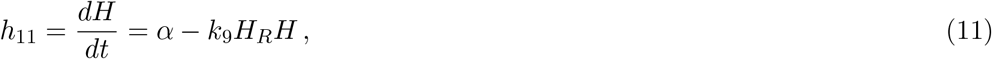

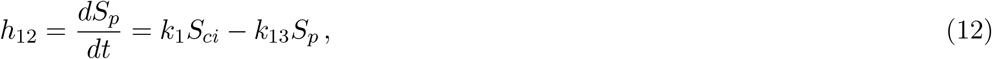

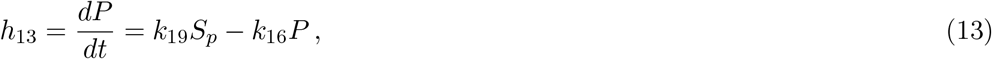

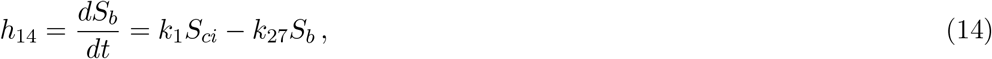

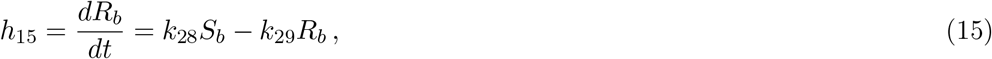

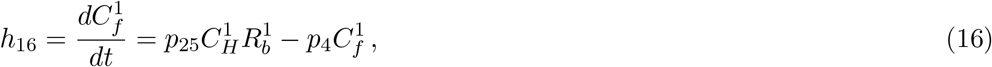

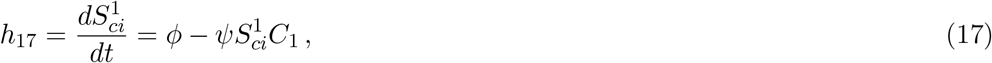

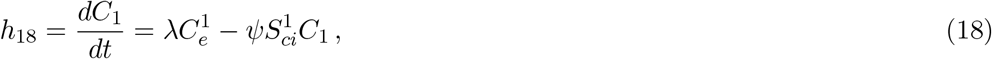

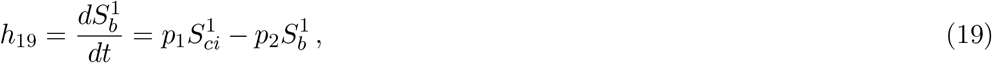

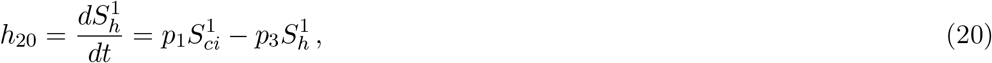

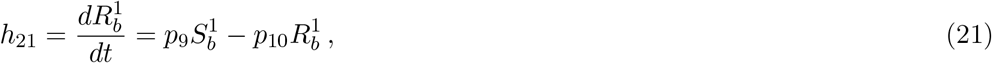

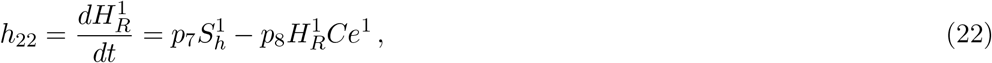

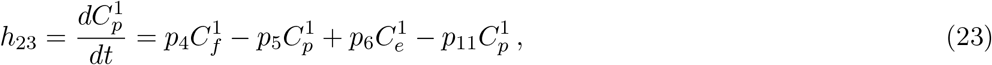

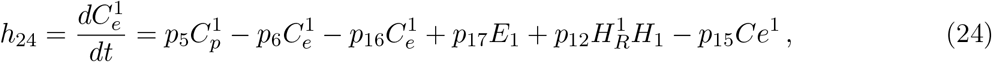

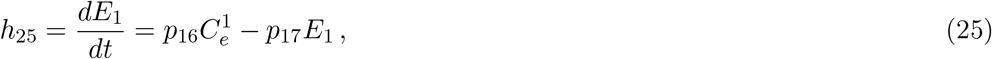

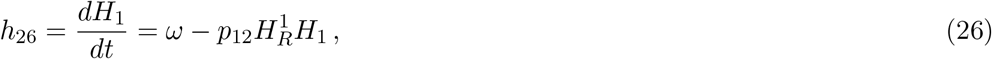

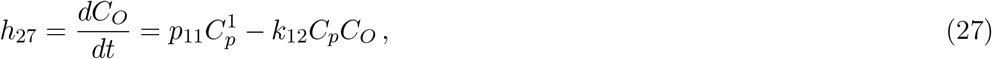

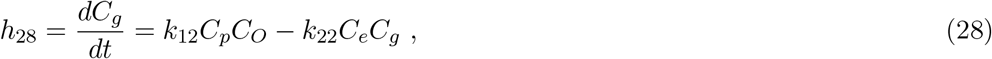

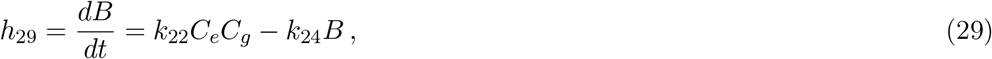

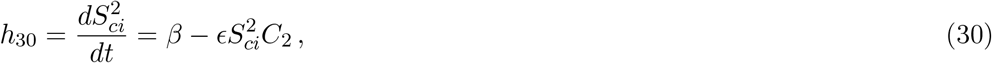

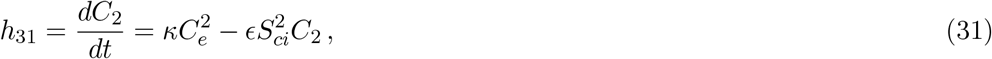

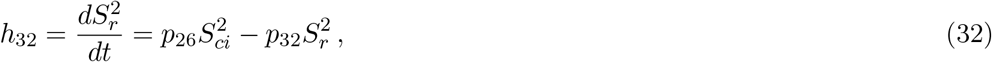

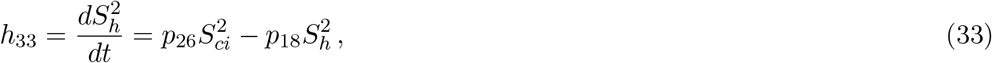

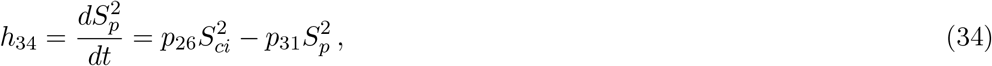

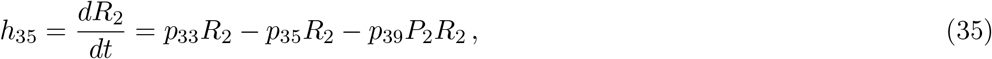

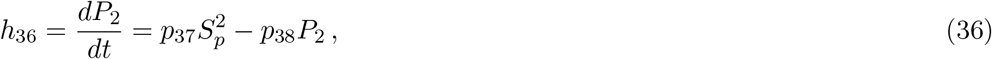

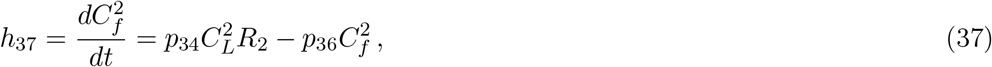

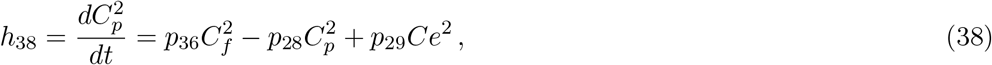

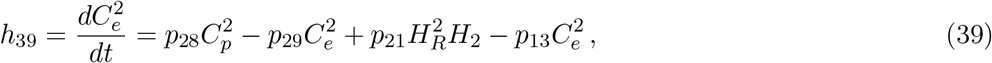

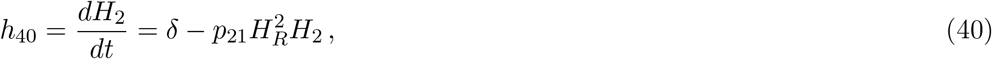

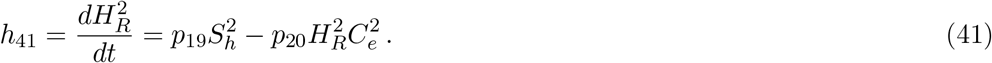

## 4 Results

### 4.1 Polynomial analysis of the retinal cholesterol CRN

Recent theoretical advances on the general principles governing RPA in arbitrarily large and complex CRNs throughout biology Araujo and Liotta (2023c, 2018); Araujo et al. (2021); Araujo and Liotta (2023b,a) now allow us to implement a systematic analysis of RPA capacity of cholesterol concentrations in the highly complex multi-compartment CRN governing cholesterol homeostasis in the retina. In particular, the Kinetic Pairing Theorem (see Theorem 1 in (Araujo and Liotta 2023c)) establishes that for every RPA-capable CRN, with interacting molecules *x*_1_, …, *x*_*n*_ and corresponding reaction rates *r*_1_, …, *r*_*n*_, it is possible to find a set of polynomials {*h*_1_, …, *h*_*n*_} ⊂ ℝ [*x*_1_, …, *x*_*n*_] such that

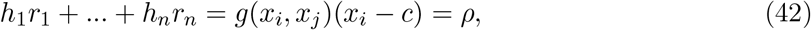

where *x*_*i*_ is any RPA-capable molecule in the CRN, with setpoint *c*, and *x*_*j*_ is any non-RPA-capable molecule in the CRN. In its simplest form, *ρ = g(x*_*i*_, *x*_*j*_*)(x*_*i*_ *− c)* constitutes the RPA polynomial of the CRN, a distinguished polynomial in two (and only two) variables, with *g*(*x*_*i*_, *x*_*j*_) its associated kinetic pairing function. Importantly, for reasons related to the underlying linearity of the CRN’s graph structure, the RPA polynomial, if it exists for the CRN under consideration, is always decomposable into a topological hierarchy of linear invariants, each obtained through only R-linear combinations of the system’s rate equations. This guarantees that the Gröbner basis computation required to test the existence of an RPA polynomial may be executed in polynomial time, comparable to Gaussian elimination.

We use this methodology here to systematically test each molecular species for RPA, using the input molecules 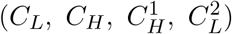 as the non-RPA-capable species in each case of the application of the Kinetic Pairing Theorem Araujo and Liotta (2023c). We have developed an open-source code for the implementation of this calculation, in the freely-available software *Singular* (https://www.singular.uni-kl.de/) within the interactive *Jupyter* environment. Our code is available at https://github.com/RonelScheepers/RPAinRetina.git. We also refer interested readers to our detailed discussion of the implementation of this systematic method for RPA-detection Scheepers and Araujo (2023) in the context of the much simpler case of single-cell-level cholesterol regulation.

This systematic method allows us to partition the species of the CRN into two disjoints sets of RPA-capable species, and non-RPA-capable species. We give a full listing of the RPA-capable species in Table 4. Our *Singular* code also automatically computes the setpoint of each RPA-capable species (also given in Table 4), as well as the combination of polynomials in the set *h*_1_, …., *h*_*n*_ (see Eq. 42) required to project the reaction rates onto the RPA polynomial. For instance, the linear combination of rate equations that identify the RPA polynomial within the steady state ideal for the ApoB–lipoprotein particle (*B*), the particle suggested by Curcio *et al*. (Curcio et al. 2011) to be the key contributor of cholesterol in drusen, is revealed by the *lift* function in *Singular* to be:

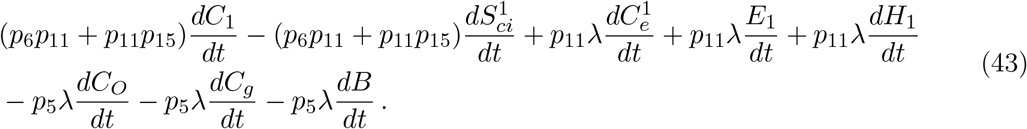

**Table 4:** RPA–capable species and their setpoints.

|  |
| --- |
| $C_e = \frac{\mu}{\theta}$ |
| $C_p = \frac{k_6 p_5 \mu \lambda + k_{21} p_5 \mu \lambda - p_5 \theta \alpha \lambda + p_6 p_{11} \theta \phi + p_{11} p_{15} \theta \phi - p_{11} \theta \lambda \omega}{k_5 p_5 \theta \lambda}$ |
| $E = \frac{k_7 \mu}{k_8 \theta}$ |
| $C_f = \frac{k_{21} p_5 \mu \lambda - p_5 \theta \alpha \lambda + 2(p_6) p_{11} \theta \phi + 2(p_{11}) p_{15} \theta \phi + 2(p_{11}) \theta \lambda \omega}{k_4 p_5 \theta \lambda}$ |
| $C_g = \frac{p_6 p_{11} \theta \phi + p_{11} p_{15} \theta \phi - p_{11} \theta \lambda \omega}{k_{22} p_5 \mu \lambda}$ |
| $B = \frac{p_6 p_{11} \phi + p_{11} p_{15} \phi - p_{11} \lambda \omega}{k_{24} p_5 \lambda}$ |
| $C_e^1 = \frac{\phi}{\lambda}$ |
| $C_p^1 = \frac{p_6 \phi + p_{15} \phi - l \lambda \omega}{p_5 \lambda}$ |
| $C_f^1 = \frac{p_5 p_{15} \phi - p_5 \lambda \omega + p_6 p_{11} \phi + p_{11} p_{15} \phi - p_{11} \lambda \omega}{p_4 p_5 \lambda}$ |
| $E_1 = \frac{p_{16} \phi}{p_{17} \lambda}$ |
| $C_O = \frac{k_5 p_6 p_{11} \theta \phi + k_5 p_{11} p_{15} \theta \phi - k_5 p_{11} \theta \lambda \omega}{k_6 k_{12} p_5 \mu \lambda + k_{12} k_{21} p_5 \mu \lambda - k_{12} p_5 \theta \alpha \lambda + k_{12} p_6 p_{11} \theta \phi + k_{12} p_{11} p_{15} \theta \phi - k_{12} p_{11} \theta \lambda \omega}$ |
| $C_e^2 = \frac{\beta}{\kappa}$ |
| $C_p^2 = \frac{p_{13} \beta + p_{29} \beta - \kappa \delta}{p_{28} \kappa}$ |
| $C_f^2 = \frac{p_{13} \beta - \kappa \delta}{p_{36} \kappa}$ |

In order to identify which molecules are directly regulated by an RPA-conferring control mechanism, and which inherit the RPA property indirectly as a result of their topological relationship to the molecules under direct homeostatic control, we have undertaken a detailed graphical analysis of the CRN. To provide context for this graphical approach, we note that recent research has focused on special cases of CRNs with special simplifying features in their interaction structures, including weakly reversible (WR) and deficiency zero (DZ) networks Feinberg (2019). However, large and complex CRNs found in real-world biological systems rarely fall into these special categories Hernandez et al. (2023). In particular, the large CRN we present here for the regulation of cholesterol in the retina eludes such simple categories. Nevertheless, a directed graph structure representing the CRN can be constructed, organising the reactions into connected components (known as ‘linkage classes’ in chemical reaction network theory Feinberg (2019)) and then decomposed into algebraically independent subnetworks (see Supplementary Information of Araujo and Liotta (2023c) for a detailed discussion of this rationale). By partitioning the CRN reactions into algebraically independent (i.e. rank additive) subsets, the steady states of the molecules within the associated subnetworks can be determined independently from their complement in the CRN. Within the context of RPA, network reactions that contribute to the network’s RPA capacity can be distinguished from reactions that play no role in the RPA capacity of the CRN Araujo and Liotta (2023c) using such decompositions.

Here we provide open-source code to accomplish the systematic decomposition of the retinal cholesterol CRN into the finest partition of independent subnetworks (see Fig. 4). This new code adapts the Matlab function developed in Hernandez et al. (2023) into a freely-available *Julia* implementation which additionally exploits the functionality of the symbolic modelling Julia package, *Catalyst*. Our Julia code is available here: https://github.com/noa-l-levi/Independent_decompositions_with_catalyst.

**Figure 4.**
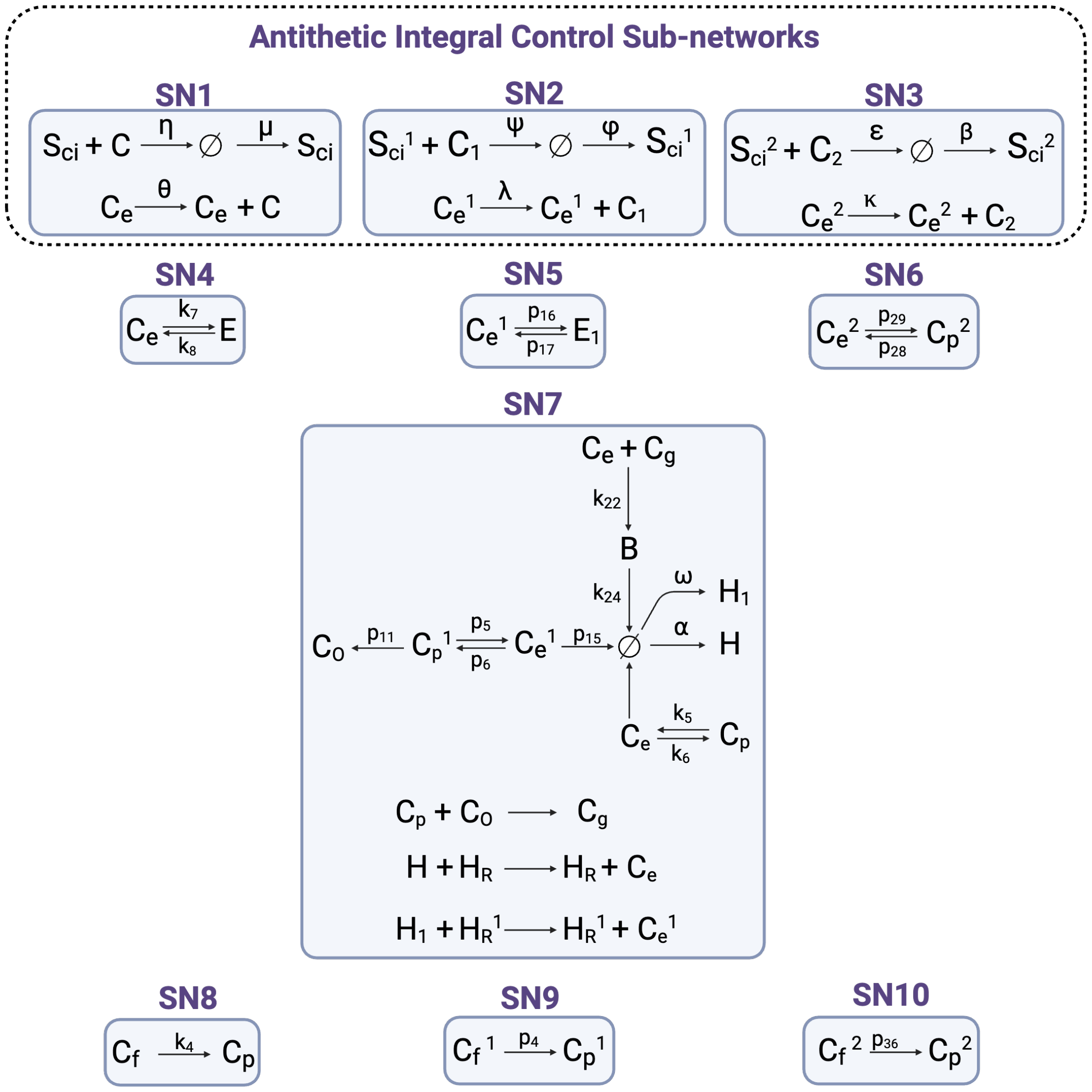
Independent Subnetworks responsible for imposing RPA on cholesterol concentrations in the human retina. (Created with BioRender.com)

Fig. 4 depicts the ten independent subnetworks that are responsible for conferring RPA on the species given in Table 4, as revealed by our Julia code, providing a set of independent graph structures that can confirm the RPA capacity of these species. All remaining reactions (i.e. those not featured in Fig. 4) are independent of the depicted reactions, and therefore do not contribute to the homeostatic regulation of cholesterol in this network.

As shown in Fig. 4, our analysis reveals the remarkable finding that there exist *three* independent antithetic integral controllers in the CRN (SN1, SN2 and SN3), each imposing RPA on 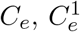 and 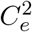, respectively. Each of these independent subnetworks has a deficiency of one, and imposes RPA on the noted species by the Shinar-Feinberg theorem Shinar and Feinberg (2010) due to the presence of the ‘null’ complex (∅) in the associated subnetwork. (We refer interested readers to a detailed discussion of these graph-theoretic technicalities in the Supplementary Information of Araujo and Liotta (2023c).) Thus, the molecules 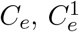 and 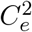 are under the direct control of the noted RPA-conferring mechanism.

By contrast, the remaining RPA-capable molecules inherit the RPA property from 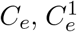 and 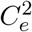 through their topological relationships to these three, via additional independent graph structures (SN4 to SN10 in Fig. 4). In particular, SN4, SN5 and SN6 are all deficiency zero, and comprise a single terminal strong linkage class, allowing E to inherit the RPA property from *C*_*e*_, *E*_1_ from 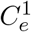 and 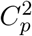 from 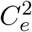. SN7 has a deficiency of three. The mass-action equations derived from this independent subset can be analysed using our above-noted *Singular* code, which confirms that the RPA property is transferred to the molecules 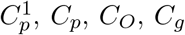and *B* from their graph relationships to *C*_*e*_ and 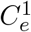. Finally, the independent subnetworks SN8, SN9 and S10 allow the transfer of RPA from *C*_*p*_ to 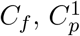 to 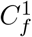 and 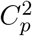 to 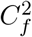. The RPA-transferring action of each of these subnetworks is independent of the actions of all other subnetworks of the CRN.

We confirm the RPA property for all above-mentioned RPA-capable molecules (as listed in Table 4) through numerical simulations of Equations (1) – (41) using a range of step functions for the input molecules (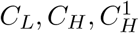 and 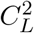), and a range of different parameter sets. We provide a representative example in Fig. 5 which illustrates the return to the expected setpoint for each RPA-capable molecule. (Parameters and initial conditions for the solutions depicted in Fig 5 are: *p*_1_ = 0.5, *p*_2_ = 16, *p*_3_ = 15, *p*_4_ = 20, *p*_5_ = 3, *p*_6_ = 2, *p*_7_ = *p*_9_ = *p*_13_ = *p*_15_ = *p*_18_ = *p*_19_ = *p*_26_ = *p*_31_ = *p*_32_ = *p*_35_ = 1, *p*_8_ = *p*_12_ = *p*_20_ = *p*_25_ = 10, *p*_10_ = 3, *p*_11_ = 0.9, *p*_16_ = 30, *p*_17_ = 20, *p*_21_ = 11, *p*_28_ = 6, *p*_29_ = 55, *p*_33_ = 20, *p*_34_ = 0.7, *p*_36_ = 0.09, *p*_37_ = *p*_38_ = 10, *p*_39_ = 2, *k*_1_ = 2, *k*_2_ = 3, *k*_3_ = *k*_6_ *= k*_11_ *= k*_*27*_ *= 1, k*_*4*_ *= 7, k*_*5*_ *= 50, k*_*7*_ *= 20, k*_*8*_ *= k*_*13*_ *= 0*.*1, k*_*9*_ *= 0*.*6, k*_*10*_ *= 1*.*2, k*_*12*_ *= k*_*19*_ *= k*_*21*_ *=* 10, *k*_*14*_ *= 0*.*5, k*_*15*_ *= 300, k*_*16*_ *= 46, k*_*17*_ *= 40, k*_*18*_ *= 20, k*_*22*_ *= 25, k*_*24*_ *= 2, k*_*26*_ *= 30, k*_*28*_ *= 40, k*_*29*_ *= 5, µ =* 2, *η* = 10, *α* = 2, *ϕ* = 50, *λ* = 40, *ω* = 1.5, *κ* = 20, *ψ* = 13, *ϵ* = 15, *δ* = 5, *θ* = 5, *β* = 10, 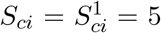, *C* = *C*_1_ = *C*_2_ = 10, 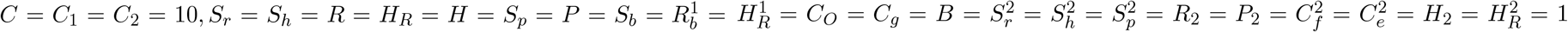, *S*_*h*_ = 2, *C*_*f*_ = *C*_*p*_ = 50, 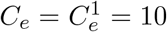, *E* = *E*_1_ = 5, *S*_*p*_ = 0.1, *R*_*b*_ = 25, 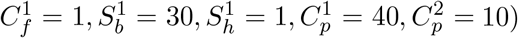.

**Figure 5.**
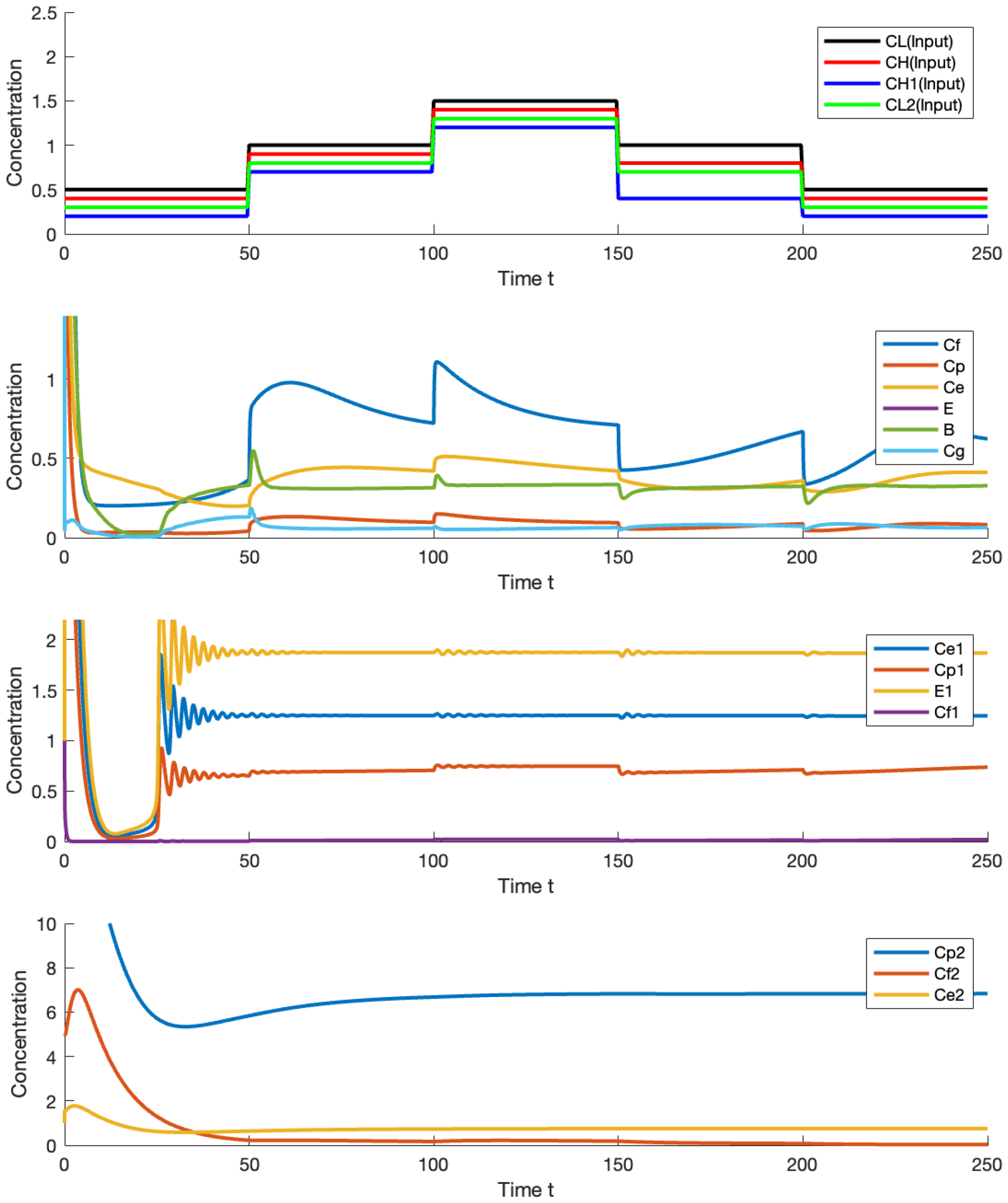
Numerical simulation of Equations (1) - (41) in response to the noted step changes in 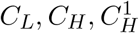 and 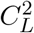. Only species exhibiting RPA are depicted here.

## 5 Discussion

Within the vertebrate retina, cholesterol is believed to exhibit a stringent form of homeostatic regulation, and thus robust perfect adaptation (RPA), which is of central importance to ensuring its proper structure and function. However, the precise molecular mechanisms underlying such robust homeostatic control have remained elusive until now. This lack of fundamental understanding of retinal cholesterol homeostasis may be attributed primarily to the complex and intricate nature of the signalling network governing cholesterol regulation, involving several dozen distinct molecular forms, compared to the relative simplicity of most documented RPA-capable CRNs. But recent technical developments in the mathematical study of universal RPA-capable CRN structures Araujo and Liotta (2023c, 2018, 2023b,a); Jeynes-Smith and Araujo (2021, 2023) have now opened up new opportunities to understand homeostatic control in complex networks such as the multi-compartment tissue-level regulation of cholesterol concentration of the human retina.

Our previous analysis of cholesterol regulation at the single-cell level Scheepers and Araujo (2023) demonstrated that an antithetic integral contoller is responsible for the tight homeostatic control of the concentration of *intracellular* cholesterol. Here, by contrast, our multi-compartment CRN representation of cholesterol regulation within the human retina, involving three connected tissue compartments of the RPE, a Müller cell layer and a photoreceptor layer, reveals a more complex picture of the homoeostatic control of cholesterol in this tissue-level setting. Nevertheless, one core feature is preserved from the simpler cholesterol regulatory CRN at the cellular level: an antithetic controller structure remains the driver mechanism for cholesterol homeostasis in several molecular forms, even in this highly complex multi-compartment CRN. In fact, in the retina, there are *three* such antithetic controllers, and each of these operates independently of the other two. Thus, from a topological viewpoint, retinal cholesterol regulation relies upon three independent *Opposer modules*, and not a *three-node opposing set* Araujo and Liotta (2018, 2023c,b,a) Moreover, these independent antithetic controllers confer RPA on endoplasmic reticulum (ER) cholesterol within their respective compartments - the RPA, the Müller cells, and the photoreceptor cells. The independence of these three distinct controllers means that any potential biochemical disruption to one mechanism (e.g. stemming from an age-related modification to the CRN network structure, or some other disease-related mutation or genomic modification) does not disturb any of the other RPA-conferring mechanisms in the retina.

From the tightly controlled ER cholesterol concentrations, RPA is transmitted via additional independent chemical reaction structures to a range of other lipid concentrations throughout the retina. Of these additional RPA transferring mechanisms, the RPE contains the most complex RPA-conferring substructure: a defiency-three subnetwork which transmits the RPA property to the ApoB–lipoprotein particle (*B*). Interestingly this is precisely the molecule suggested by Curcio *et al*. (Curcio et al. 2011) to be the key contributor of cholesterol accumulation in drusen - an important driver in the development of macular degeneration. Importantly, homeostasis of the ApoB–lipoprotein particle (*B*) is regulated by a more complex mechanism than antithetic integral control. This newly-identified homeostatic mechanism may now offer insights to the scientific community in examining potential mechanisms for perturbation to this robust cholesterol homeostasis in the context of diseases such as age-related macular degeneration (AMD).

The mathematical framework we present here makes clear that the constitutive production of the SREBP/SCAP/Insig complex (*S*_*ci*_), as well as the sensing of *active* cholesterol molecules (*C*_*i*_) by this complex, is fundamental to the robust homeostatic regulation of cholesterol at the cellular level and ultimately throughout the retina. Of the main sterol regulatory-element binding proteins, SREBP2 regulates the transcription of genes involved in cholesterol metabolism. A recent review by Shimano and Sato Shimano and Sato (2017) points to several studies that have investigated the pathophysiological influence of SREBP2 at the cell, tissue, organs and organism level. For example, it is hypothesised that the up–regulation of cholesterol biosynthesis in many cancers may be due to mutations in the gene transcription of SREBP2 (Zahra Bathaie et al. 2017), while it has also been shown that the mammalian target of rapamycin complex 2 protein Kinase B (TORC2–AKT) signalling may cancel the activation of SREBPs. This dysregulation might contribute to insulin resistance and diabetes mellitus Williams et al. (2013). Furthermore, when SREBP2 is overexpressed in pancreatic *β*–cells of mice, it induces *β*–cell death, thereby disrupting glucose tolerance and potentially leading to diabetes mellitus Ishikawa et al. (2008). Given SREBP2’s role in cholesterol synthesis, novel therapeutic strategies targeting it hold promise for treating cancers with abnormal cholesterol metabolism Xue et al. (2020). While evidence exists for the effects of disrupted SREBP2-regulated cholesterol metabolism in vertebrate and mammalian cells Shimano and Sato (2017), to our knowledge, no similar studies on the impact of SREBP2 disruption in mammalian retinal cells exist currently.

## Author Contributions

RS: conceptualisation, formal analysis, methodology, software„ visualisation, writing - original draft preparation. NLL: methodology, software, formal analysis; RPA: conceptualisation, methodology, writing - review and editing, supervision.

## Conpeting Interests

The authors have no competing interests to declare.

## Funding

Robyn P. Araujo is the recipient of an Australian Research Council Future Fellowship (project number FT190100645) funded by the Australian Government

## Notes

### Competing Interest Statement

The authors have declared no competing interest.

